# Feeding niches drive specialist herbivore responses of conspecific insects to plant defenses in biocontrol

**DOI:** 10.1101/2025.01.29.635475

**Authors:** Jasleen Kaur, Lucia Navia, Emily Kraus, Octavio Menocal, Philip G. Hahn

## Abstract

1. Biological invasions threaten global ecosystems and economies, prompting the need for sustainable management approaches such as releasing natural enemies from the native range (i.e., classical biological control; CBC). While the release of multiple natural enemies can enhance the effectiveness of CBC, interactions between these agents and their host plants’ defenses are poorly understood.
2. This study explored the interactions between two congeneric specialist herbivores introduced for the control of *Dioscorea bulbifera* (air potato) in Florida, USA: *Lilioceris cheni* (Coleoptera: Chrysomelidae), which feeds primarily on leaves and *L. egena*, which feeds primarily on the aerial reproductive structures (i.e., bulbils). Specifically, we investigated whether foliar herbivory by *L. cheni* or induced defenses by jasmonic acid (JA) or salicylic acid (SA) influenced *L. egena*’s feeding behavior and performance on bulbils.
3. We conducted feeding assays to measure bulbil consumption and its relationship to pupal mass and adult eclosion as indicators of beetle performance. Additionally, we performed dose-dependent feeding assays to test how varying concentrations of saponins, a key defense compound in air potato, influenced feeding and survival of *L. cheni* and *L. egena*.
4. Our results indicated no significant differences in bulbil feeding by *L. egena* across treatments; however, SA-treated plants produced significantly heavier pupae and adults compared to controls. Dose-response assays revealed non-linear, hump-shaped, feeding and survival patterns in *L. cheni*, while *L. egena* experienced reduced feeding and survival with increasing saponin concentrations.
5. These findings suggest that the two beetles respond differently to plant defenses while occupying distinct feeding niches, likely allowing for additive impacts on *D. bulbifera*. This study provides insights into the role of plant defenses in CBC and informs strategies to optimize biocontrol programs for invasive species management.

## Introduction

Invasive plants pose a threat to biodiversity by disrupting ecosystem functions like nutrient cycling, fire regimes, and hydrology, thereby impacting the local flora and fauna (Mack *et al*., 2000; Walsh *et al*., 2023). Classical biological control (CBC) is a popular tool to counteract these negative ecological effects of invasive species by strategically introducing natural enemies from the native range of the invasive pest (van Driesche *et al*., 2010; van Driesche, 2012; Lake & Minteer, 2018). In their native range, plants are often attacked by multiple natural enemies (Schoonhoven *et al*., 2005). Consequently, a logical consideration for CBC is to exploit potential synergies arising from the simultaneous presence of these natural enemies. Moreover, there is evidence supporting that the release of multiple species of biological control agents is more effective in reducing pest populations than the release of a single species (Stiling & Cornelissen, 2005). This increased effectiveness can be attributed to both the cumulative damage inflicted by individual species, in line with the cumulative stress hypothesis, and the increased likelihood of introducing a successful agent, as predicted by the lottery model (Stephens *et al*., 2013). The cumulative stress hypothesis posits that the augmentation of plant suppression occurs by accumulating damage from multiple species over time (Swope & Parker, 2010), whereas the lottery model suggests that introducing a greater number of biological control agents increases the probability of introducing a successful one (Denoth *et al*., 2002). However, addition of multiple agents also adds to the ecological complexity, which may increase the odds of deleterious outcomes (Impson *et al*., 2008). Nevertheless, the efficacy of deploying multiple natural enemies remains a subject of debate among researchers, with particular concerns centering on potential interference between these agents that could compromise the overall success of biocontrol initiatives (Stiling & Cornelissen, 2005; Pearson *et al*., 2022).

Compatibility of introduced agents is essential to ensure their collective efficacy as their interaction can result in potential synergistic, antagonistic, or additive effects, when compared to their individual impact. For instance, competition is the most likely outcome when multiple natural enemies share a single host plant (Kaplan & Denno, 2007). The competitive exclusion principle posits that if two coexisting populations of different species, with varying population growth rates, occupy the same ecological niche, the competitively dominant species will replace the other (Hardin, 1960; Cushing *et al*., 2004; Johnson & Bronstein, 2019). However, competing species can coexist through niche differentiation, involving mechanisms like spatial or temporal resource partitioning or partitioning along a resource spectrum (Chase & Leibold, 2003). Niche differentiation in the form of feeding on different plant structures (i.e., leaves and fruits) could result in coexistence of the herbivores as biocontrol agents and add to the cumulative stress for the plant. For instance, literature suggests that the coexistence of aboveground (*Cyphocleonus achates* Fahraeus) and belowground (*Larinus minutus* Gyllenhal) herbivores had additive negative effects on the performance of spotted knapweed (*Centaurea stoebe* L.), resulting in reduced growth and reproduction success over time. This indicates that such differentiation can lead to cumulative negative effects on the host plant (Knochel *et al*., 2010; Stephens & Myers, 2014).

Multiple biological control agents can also interact indirectly, through plant-mediated effects such as induced defenses (Milbrath & Nechols, 2004). There can be antagonistic interactions, where the combined herbivory effect on the plant may fall short of the impact of individual species. This may arise when attack by one enemy induces plant defenses or alters plant phenology that reduces host susceptibility to the second enemy (Bezemer *et al*., 2003; Milbrath & Nechols, 2004; Stephens *et al*., 2013). For example, leaf-chewing herbivores, such as *Plutella xylostella* L. (Lepidoptera: Plutellidae) and *Pieris brassicae* L. (Lepidoptera: Pieridae), induce plant defenses in *Brassica oleracea* L. that negatively affect the performance of root-feeding *Delia radicum* L. (Diptera: Anthomyiidae) larvae (Soler *et al*., 2012; Karssemeijer *et al*., 2022). Alternatively, one natural enemy may facilitate another resulting in their combined herbivory exceeding their individual effects. This is possible when attack by the first enemy alters the nutritional profile or plant phenology, rendering the host plant more susceptible to the second enemy (Masters *et al*., 2001; Johnson *et al*., 2009; Stephens *et al*., 2013). These nuanced dynamics underscore the importance of understanding and predicting the interactions among multiple natural enemies for the effective implementation of biocontrol strategies as this will facilitate the species selection for biological control introductions. Therefore, to create a comprehensive approach, it is crucial to address the ecological and economic factors, satisfying the need for well-informed strategies in classical biological control.

Given that numerous CBC agents have been released to manage individual invasive species (Schwarzländer *et al*., 2018) there is significant potential for CBC agents to interact in unexpected ways. One such system to investigate these potential interactions involves the interactions between specialist herbivores as natural enemies, *Lilioceris cheni* Gressit & Kimoto and *Lilioceris egena* (Weise) of the highly invasive plant air potato (*Dioscorea bulbifera* L.) in the southeastern United States (Croxton *et al*., 2011; Manrique *et al*., 2023). Both *L. cheni* and *L. egena* have been approved for release and they can feed on different structures of the plant—leaves and bulbils (Center *et al*., 2013; Overholt *et al*., 2016; Smith *et al*., 2018). However, *L. cheni* is primarily a leaf feeder whereas *L. egena* prefers to feed primarily on bulbils, the reproductive structures that enable the plant to reproduce asexually and overwinter in the soil (Dray *et al*., 2023). This specialization is expected to reduce competition between them as each herbivore occupies a distinct ecological niche. As a result, their combined impacts are expected to work additively, enhancing biocontrol effectiveness, as predicted by the cumulative stress hypothesis.

Multiple studies suggest that when chewing insects attack, the Jasmonic acid (JA) pathway is activated, enhancing the plant’s defenses by inducing the production of compounds that deter their herbivore feeding and performance, while Salicylic acid (SA) pathways respond primarily to pathogen attacks and piercing insects like aphids (Howe & Jander, 2008; Thaler *et al*., 2012). Additionally, air potato is rich in plant defense compounds such as steroidal saponins, particularly diosgenin (Ghosh *et al*., 2013; Badole & Dighe, 2019; Raina & Misra, 2020). Saponins can negatively affect insect physiology by disrupting cellular membranes and digestive processes in herbivores. This inducibility and the inherent toxicity of saponins underscore the defense mechanisms of *D. bulbifera*, which play a crucial role in its interactions with its natural enemies. Previous findings revealed that induction by JA and *L. cheni* feeding damage led to an increase in the production of phytochemicals, specifically saponins, which subsequently enhanced further feeding by the same beetle (Kaur *et al*., 2023). The *Lilioceris* beetles, are highly specialized on their host plant (Center *et al*., 2013; Lake *et al*., 2015) as well as colored bright red (aposematic). Therefore, these beetles may use sequestration as a counter-defense mechanism (Ernst, 2005). Sequestering specialists can benefit from low to intermediate levels of plant toxins, though their performance may decline at higher toxin levels. Conversely, specialists that do not sequester chemicals may not experience significant effects from plant defenses unless toxin levels are high enough to be detrimental. In contrast, generalist insects typically exhibit a dose-dependent, linear negative response to plant defenses (Ali & Agrawal, 2012).

Based on this understanding and previous observation, in this study we used feeding assays to examine whether the defense induction in air potato using the following treatments - prior foliar damage by *L. cheni*, JA, SA and water as control spray influences the subsequent feeding levels of *L. egena* on the bulbils that are produced later in the season. We hypothesized that 1) *L. egena* larvae would have higher feeding and performance (i.e., Pupal mass and adult eclosion) on bulbils derived from JA-treated plants and plants with previous leaf damage from *L. cheni* beetles (as these may be induced in the bulbils upon *L. cheni* herbivory on the foliage of air potato earlier in the growing season) compared to SA and control treatment. We also examined how bulbil consumption by *L. egena* larvae relate to their performance on the host plant as a measure of food quality.

Additionally, we investigated the dose dependent effect of the saponins on both *L. cheni* and *L. egena* beetles’ feeding and development through lab-based feeding assays on artificial diet. Therefore, we hypothesized that 2) both *L. cheni* and *L. egena* exhibit positive feeding responses, weight gain, and frass production to saponin concentrations ranging from low to intermediate concentrations and then decline at high concentrations forming a hump shaped curve, in concordance with expectations for sequestering specialists (Ali & Agrawal, 2012). The outcomes of this research provide valuable insights into optimizing strategies for the effective management of air potato.

## Material and methods

### Study insects

*Lilioceris cheni* beetles as well as *L. egena* larvae and adult beetles were used for this study to investigate their interactions and effects on air potato. The beetles were reared under laboratory conditions, providing a stable and accessible source for experimental work. For this project, the Chinese biotype was chosen due to its availability from the Florida Department of Agriculture and Consumer Services - Division of Plant Industry (FDACS-DPI) and its demonstrated performance under controlled rearing conditions. This consistency ensures reliable results when investigating its interactions with *L. egena* and cumulative effects on air potato, aligning with the study’s objectives.

Both adult beetle species and freshly emerged larvae of *L. egena* were obtained from FDACS-DPI in Gainesville, FL, USA. These insects were mass reared under controlled conditions according to standard procedures, kept at 27 ± 2 °C, 65 ± 10% RH, and 14:10 Light: Dark cycle (Kraus *et al*., 2022; Murray *et al*., 2025).

### Bulbil feeding experiment- *L. egena*

#### Plant rearing and experimental design

Plants in this study were grown from bulbils (67 – 277 g) collected in Florida by FDACS-DPI and planted in March 2023. Bulbils were planted individually in 4 L pots filled with a soil potting medium (Jolly Gardener, Pro-Line C/25 Growing Mix) under natural light conditions in greenhouse (Temperature 26 ± 5 °C, RH 75%). Each pot was equipped with four 1.5 m tall bamboo stakes and a cylindrical cage made of tule mesh material (Item # 1825165, Jo-Ann Stores, Inc.). The mesh was secured to the pot using rubber bands at the base and tied at the top with a zip tie. To maintain optimal moisture levels, each pot was watered 1-3 times per week.

Plants were grown for three months and one of the four treatments (n= 20 per treatment) were randomly assigned to them: 1) sprayed with methyl jasmonate (MeJA; conjugate of JA) at a concentration of 5 mM, 2) sprayed with salicylic acid (SA), 3) *L. cheni* herbivory was induced by releasing three freshly emerged beetles until the plants were damaged to ∼60 percent, 4) a control group where plants were sprayed solely with water. Each plant was sprayed with 3.5 ml of the assigned treatment solution. Solutions for MeJA and SA acid were prepared following the methodologies outlined in previous studies (Khodary, 2004; Kraus & Stout, 2019; Kaur *et al*., 2023). The treatments were repeated approximately one month after the initial treatment and a month before harvesting the bulbils. The repeated application of the treatments was to ensure the plants continued to induce their defensive systems. Additionally, it is important to note that plants in the *L. cheni* herbivory treatment had accidental beetle oviposition that resulted in a few clutches of freshly emerged larvae. However, they were immediately removed during daily monitoring operation on the plants. Finally, the bulbils were harvested from the plants upon growing until seven months for conducting the feeding assay.

### Experimental assay and data collection

We conducted this experiment in three waves (i.e., temporal blocks), ensuring that each wave comprised bulbils from all treatments and of comparable sizes as the bulbils displayed variations in both size and quantity across all plants. Bulbils used in this experiment ranged from 0.87 to 16.81 g. To ensure a judicious use of bulbil material, and to have the amount of feeding material similar across all samples, the bulbils were sliced into similar sections. Therefore, a total of 99 bulbil slices were used across all treatments, comprising 14 from JA, 27 from SA, 25 from *L. cheni* herbivory, and 33 from the control treatments. Each sliced bulbil sample received one neonate larva to test the effects of each treatment on the larvae feeding level. The amount of bulbil material provided to each larva was large enough to ensure it had enough food for the duration for the experiment.

The samples were individually placed in separate 60 ml clear plastic deli cups, covered with a lid and placed at room temperature. Larval frass from the entry site of the larva in the bulbil was collected daily and stored in a 2 ml Eppendorf tube. The frass collection from each larva was accumulated throughout the experiment to determine total weight of frass produced by the larva. Larvae were allowed to feed until pupation. Upon reaching full maturity, larvae exited and dropped into vermiculite (2 cm deep layer) that was placed beneath the bulbil in the deli cup for pupation. Mortality was recorded for the duration of the experiment.. In total, the experiment lasted for 12-16 days, from the placing of freshly emerged larvae on the bulbil until their pupation.

To quantify larval damage after pupation, we used a modification of the protocol designed by Dray & Goldstein, (2019). Each bulbil sample was further sliced into small pieces at a thickness of 0.8 mm using a vegetable slicer. Beeswax was placed in a 10 ml beaker and heated in a water bath. Then, the molten beeswax was dispensed into cavities formed upon insect feeding on each bulbil slice using a 1000 ml pipette, which was removed upon hardening. Wax from all slices of a single bulbil was pooled and weighed as a proxy of the amount of bulbil material consumed. Additionally, we measured pupal and adult weight.

### *Data analysis:* Bulbil feeding experiment- *L. egena*

Data were analyzed in R (Version 1.3.1093). To investigate the performance of *L. egena* beetles on bulbils from plants subjected to prior leaf damage by *L. cheni*, applications of phytohormones (JA or SA), and control treatments, we first conducted a multivariate analysis of variance (MANOVA). This analysis utilized the lm function, to assess multiple response variables - frass weight, pupae weight, wax weight (as a proxy for beetle damage), and beetle weight gain. It employed the Wilks’ lambda statistic to test for significant mean differences among treatment groups. Thereafter, separate linear models were fitted for each response variable using the glmmTMB package (Brooks *et al*., 2017), with the treatment as fixed effect. For each response variable, model diagnostics were conducted using the simulateResiduals function from the DHARMa package (Hartig, 2020) to evaluate the residual normality and homoscedasticity.

To further explore the relationship between bulbil consumption by *L. egena* larvae and their performance on air potato plants as a measure of food quality, we fitted generalized linear mixed models (GLMMs) using the glmmTMB package. We included food consumption and treatment, as well as their interaction, using Wald χ^2^tests such that wax weight and frass weight as a proxy of food consumption were included in separate models. Both models included pupa weight (performance) as a response variable.

The Anova function from the car package (Fox & Weisberg, 2019) was used to test the significance of treatment using Wald χ^2^ tests, and post-hoc linear contrasts using Tukey-adjusted *p*-values were constructed with emmeans function in emmeans package. Subsequent pairwise comparisons among treatment levels were also conducted using the emmeans package (Lenth, 2016). We also calculated emtrends from emmeans package for factor-covariate interactions and estimates the slopes of the covariate trends at each factor level.

### Dose dependent bioassay - *L. egena* and *L. cheni*

#### Experimental assays and data collection

We conducted dose dependent feeding bioassays on artificial diets for both *L. cheni* and *L. egena.* For these experiments, an artificial diet using the commercially available Colorado potato beetle diet (Frontier Agricultural Sciences, USA) was formulated as per the company instructions with some modifications. We also added KOH and neomycin to manage the mold growth in the diet. Additionally, to make sure the specialist beetles feed on the artificial diet, air potato leaf material was also included at the minimum possible quantity (0.4 g) enough to initiate insect feeding (Kraus *et al*., 2022). Then, variable levels of diosgenin (Sigma-Aldrich #D1634-5G) —a saponin derived from air potatoes were added to create diosgenin level treatments ranging from low to high as 0, 2.5, 5, 10, 15 and 20 mg/g for *L. cheni* and 0, 15, 25, 35, 50 mg/g of *L. egena*. These levels were based on the levels of diosgenin naturally found in air potato plants (Kaur *et al*., 2023), but expanded so that we had a range of diosgenin levels slightly beyond the natural range. The detailed quantities for artificial diet formulations for both *L. cheni* and *L. egena* are provided in the supplementary file (Table S1, Table S2). While artificial diet comprised of leaf content for *L. cheni* beetles, the saponin levels were modulated by incorporating 0.4 g of bulbil powder instead of air potato leaf material. Both leaf and bulbil powders were obtained by freeze drying (Harvest Right, the USA) them for 24 hours, and subsequently milling using zirconium beads and the BeadBug homogenizer (BeadBug™ 6 microtube homogenizer, Millipore Sigma).

We conducted the experiment in two waves for *L. cheni* and three waves for *L. egena*, with each dosage treatment being replicated total of 24 times. Therefore, the experiment for *L. cheni* comprised of total 144 samples (24 × 6 treatments) across all treatments, while *L. egena* consisted of 120 samples (24 × 5 treatments) across all treatments. The setup involved 60 ml clear plastic deli cup containers with punctured lids to facilitate air circulation. A paper disc at the base of the container supported a diet piece (∼7 mm radius). All the containers were placed on a lab bench under temperature 22-24°C. The experimental adult beetles, aged 1-2 days post-pupal eclosion, were subjected to a 48-hour and 24-hour starvation period, for *L. cheni* and *L. egena* respectively, and were weighed before placement on the diet. Diet was replaced daily to prevent contamination by mold growth and daily collections of frasses were immediately stored in separate 2 ml Eppendorf tubes at −20° C, for subsequent data collection of frass weight for each beetle. The experiment concluded after 10 days of feeding. Mortality was recorded for all insects that died during the experiment.

### Data Analysis: Dose dependent bioassay - L. cheni and L. egena

Data were analyzed in R (Version 1.3.1093). For this experiment, the data was screened for outliers, leading to the exclusion of a single outlier—a *L. cheni* beetle with an unusually high frass weight entry that also died early in the experiment. To investigate the dose-dependent effects of saponins on the feeding behavior and performance of *L. cheni* and *L. egena* on the artificial diet, we analyzed the response variables—frass weight and survival—separately using Generalized Linear Models (GLMs) with the glmmTMB package (Brooks *et al*., 2017). Each beetle species was also analyzed separately (four models total). Frass weight produced by each beetle was modeled with a normal distribution, while survival (alive or dead for each beetle) was modeled with a binomial distribution. For each response variable and each species, three candidate models were constructed: a null model, a linear model, and a quadratic model. Among the models, the null model only included an intercept as the predictor variable, the linear model included the treatment (i.e., the diosgenin levels; continuous variable) as the predictor, and the quadratic model incorporated both the diosgenin dose treatment and its square as the predictor variables. The linear model tests for directional responses of the beetle to saponin concentrations while the quadratic model tests for potential unimodal patterns as predicted for sequestering specialists. The null model helps with comparison for no effect of diosgenin levels on feeding or performance of beetles.

Model comparison was based on the Akaike Information Criterion (AIC) value to compare the model’s goodness of fit. Models were selected based on the lowest AIC score, models having a delta-AIC value of less than 2 were considered the best models. Further, we used the summary function from base R to examine the model output details. Finally, we employed the r.squaredGLMM function from the MuMIn package (Barton & Barton, 2015) to assess the proportion of variance explained by the model.

## Results

### Bulbil feeding experiment- *L. egena*

The MANOVA indicated a significant effect of treatment on bulbil consumption metrics (Wilks’ Λ = 0. 744, F (df = 12, 204.01) = 2.011*, p =* 0.025), suggesting differences among the treatment groups. Subsequent results from Univariate ANOVA did not reveal any statistically significant differences in wax weight as a proxy for beetle damage across treatments (χ^2^ = 1.288, df = 3, *p =* 0.732; Figure 1a). The analysis for frass weight also showed no significant differences among treatments (χ^2^ = 3.811, df = 3, *p* = 0.283; Figure 1b). However, our results showed a significant effect of treatment on pupae weight (χ^2^ = 18.816, df = 3, *p* = 0.0003; Figure 1c). The SA treatment resulted in the highest pupae weights (mean = 143 mg ± 4.72 SE, 95% CI= 134, 153), significantly more than those on the control (mean = 116 mg ± 4.72 SE, 95% CI= 107, 124*, p=* 0003), but not significantly higher than JA treatment (mean = 127 mg ± 6.53 SE, 95% CI= 114, 140*, p=* 0.208) or *L*. *cheni* damaged beetle treatment (mean = 131 mg ± 4.72 SE, 95% CI= 121, 140*, p=* 0.252). Additionally, there was no difference found between JA and *L*. *cheni* damaged beetle treatment (*p=* 0.974), control and JA treatment (*p=* 0.454) as well as control and *L*. *cheni* damaged beetle treatment (*p=* 0.095).

**Figure 1.**
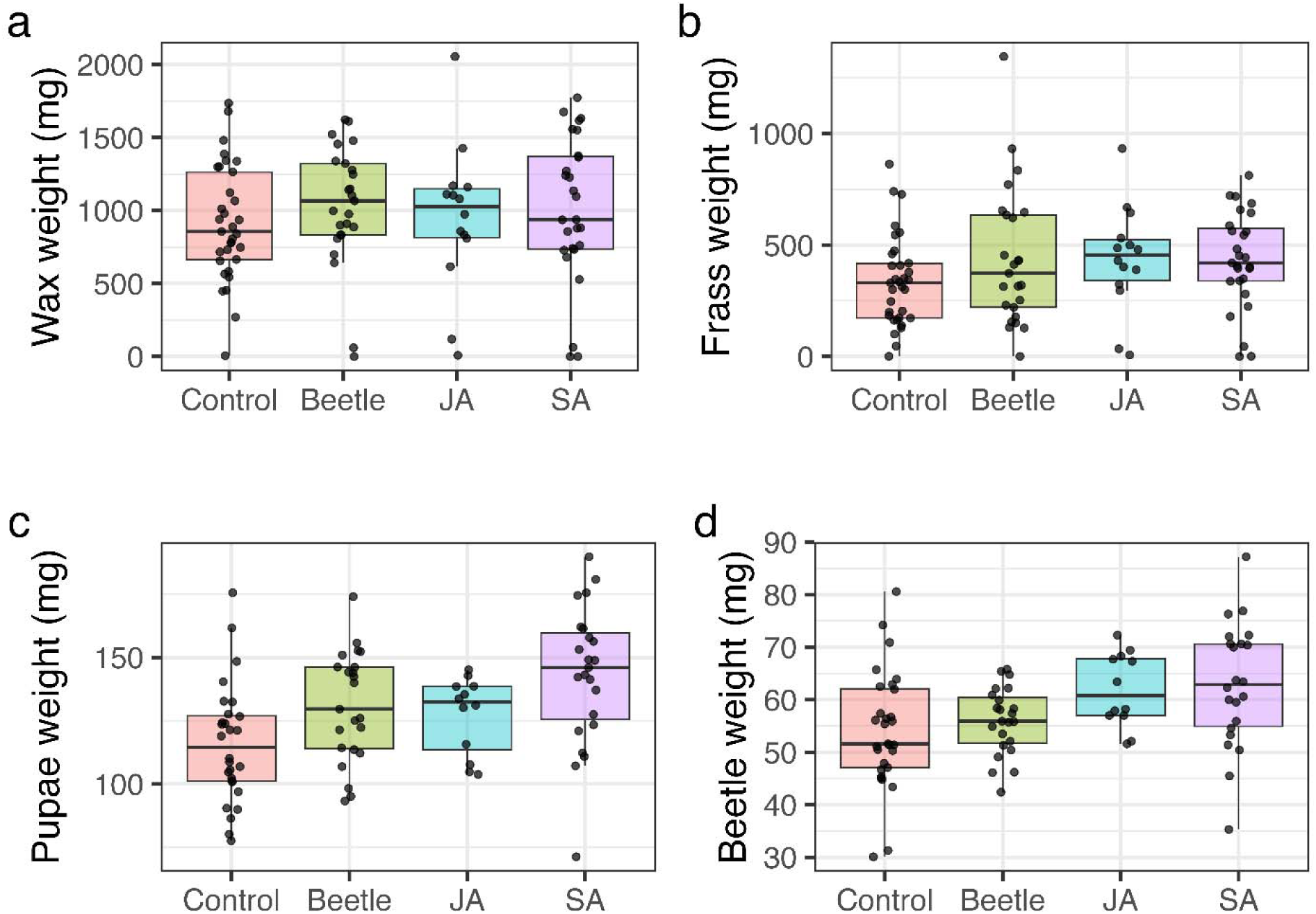
Effects of different treatments on feeding by *L. egena* on treatment bulbils. (a) Wax weight (mg) (b) Frass weight (mg) oflarvae, (c) Pupae weight (mg) (d) Beetle weight (mg) of *L. egena* across four treatment groups of induced defenses: Control, Beetle (prior herbivory by *L. cheni*), Jasmonic Acid (JA), and Salicylic Acid (SA). Each color represents distinct treatment across each panel. Points are shown for individual beetles and boxplots show medians and interquartile ranges, with whiskers representing 95^th^ quartile range.

We also found a significant effect of treatment on weight of emerging beetles (χ^2^ = 18.816, df = 3, *p* = 0.004; Figure 1d). Like pupae weight, the SA treatment resulted in the highest beetle weights (mean = 62.8 mg ± 2.05 SE, 95% CI= 58.8, 66.9), significantly more than those on the control (mean = 54.1 mg ± 1.78 SE, 95% CI= 50.5, 57.6, *p=* 0.009), but not significantly higher than JA treatment (mean = 61.8 mg ± 2.77 SE, 95% CI= 56.3, 67.4*, p=* 0.992) or *L*. *cheni* damaged beetle treatment (mean = 55.9 mg ± 2.0 SE, 95% CI= 52, 59.9*, p=* 0.084). Additionally, there was no difference found between JA and *L*. *cheni* damaged beetle treatment (*p=* 0.316), control and JA treatment (*p=* 0.094) as well as control and *L*. *cheni* damaged beetle treatment (*p=* 0.899).

We found that frass weight, a proxy of bulbil consumption, significantly predicted pupae weight (χ^2^ = 18.332, df = 1, *p* < 0.0001), and treatment also had a significant effect (χ^2^ = 13.596, df = 3, *p* = 0.003). The interaction between frass weight and treatment was not significant (χ^2^ = 2.480, df = 3, *p* = 0.479). We found a significant positive correlation between pupae weight and frass weight in control (trend = 0.0508 ± 0.0226 SE, t-ratio = 2.243, *p* = 0.0278; Figure 2a), in *L*. *cheni* damaged beetle treatment (trend = 0.0385 ± 0.0144 SE, t-ratio = 2.669, *p* = 0.0093; Figure 2b), and in the SA treatment (trend = 0.0745 ± 0.0255 SE, t-ratio = 2.921, *p* = 0.0046; Figure 2d). The JA treatment showed no significant trend (*p* = 0.7254; Figure 2c). Additionally, wax weight, significantly predicted pupae weight (χ^2^= 23.796, df = 1, *p* < 0.0001), and treatment also had a significant effect (χ^2^ = 14.058, df = 3, *p* = 0.003). Similar to the previous model, the interaction between wax weight and treatment was not significant (χ^2^ = 2.874, df = 3, *p* = 0.411). We found a significant positive correlation between pupae weight and wax weight in the SA treatment (slope = 0.04 ± 0.011 SE, t-ratio = 3.48, *p* = 0.0008) and in *L*. *cheni* damaged beetle treatment (slope = 0.0439 ± 0.0144 SE, t-ratio = 3.048, *p* = 0.0032), while the control showed a weaker but significant trend (slope = 0.0235 ± 0.0111 SE, t-ratio = 2.113, *p* = 0.0378). The JA treatment showed no significant trend (*p* = 0.3726).

**Figure 2.**
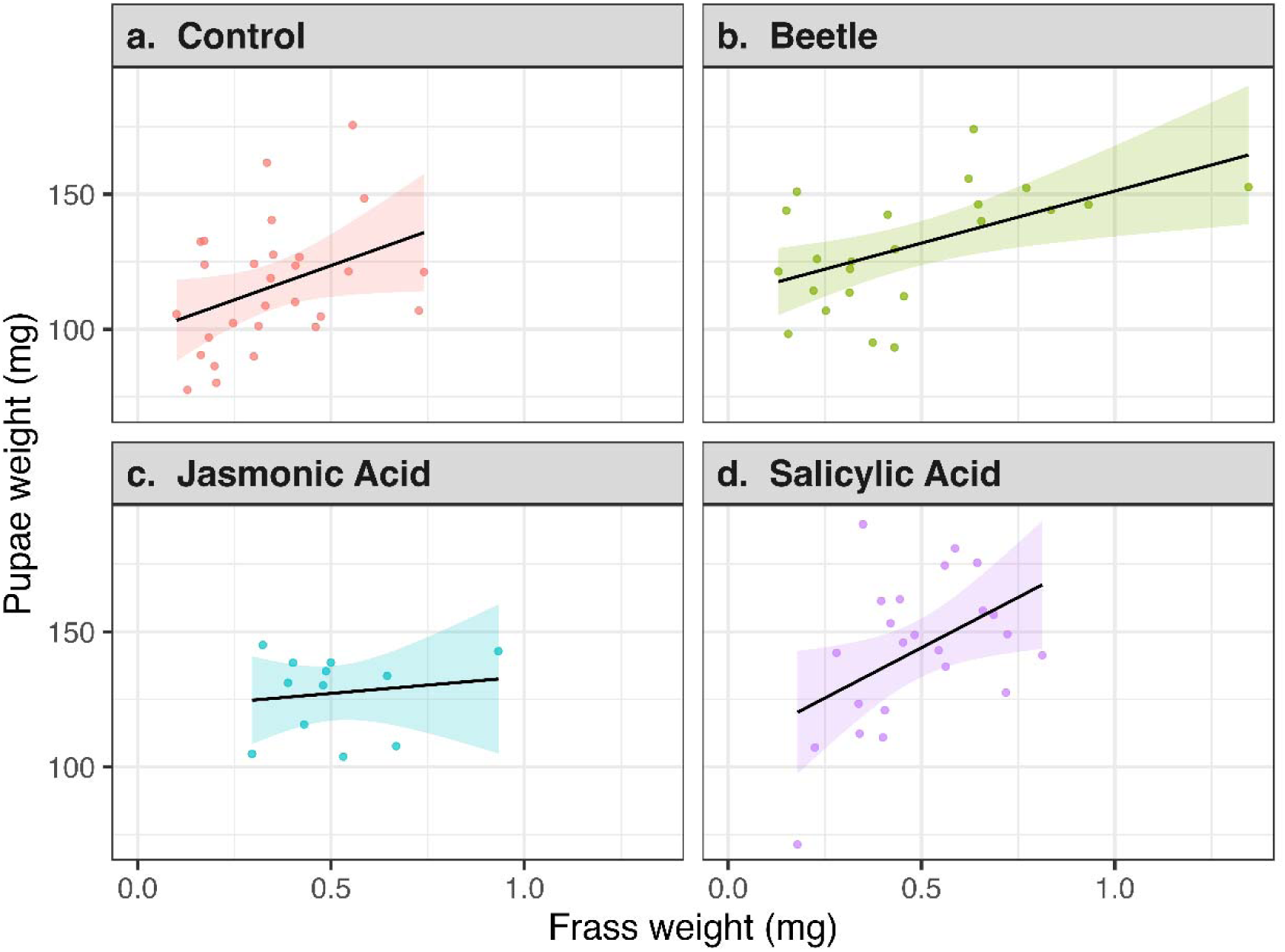
Relationship between frass weight (mg) (food consumption) and Pupae weight (mg) (insect performance) across four treatment groups of induced defenses in separate panels: Control, Beetle (prior herbivory by *L. cheni*), Jasmonic Acid (JA), and Salicylic Acid (SA). Each color represents distinct treatment in each panel. Each plot shows points for individual insects in designated treatment and a fitted curve with shaded areas represent the 95% confidence intervals.

### Dose dependent bioassay - *L. cheni* and *L. egena*

Three models were compared to test the dose-dependent effects of saponins on the frass weight of *L. cheni.* Frass weight was used as the response variable, against four different doses of diosgenin as continuous predictor variable. The quadratic model had the lowest AIC value (828.31). However, the linear model also had a competitive AIC value (829.86). Nonetheless, both models were better than the null model (AIC= 832). Subsequent evaluation revealed that the frass weight from *L. cheni* increased initially at lower to intermediate concentrations (Slope of linear term = 1.195 ± 0.503 SE, z = 2.378, *p =* 0.017; Figure 3a), and declined thereafter at higher concentrations of diosgenin (Slope of squared term = −0.045 ± 0.024 SE, z = −1.897, *p =* 0.058, *R^2^* = 0.067, Figure 3a).

**Figure 3.**
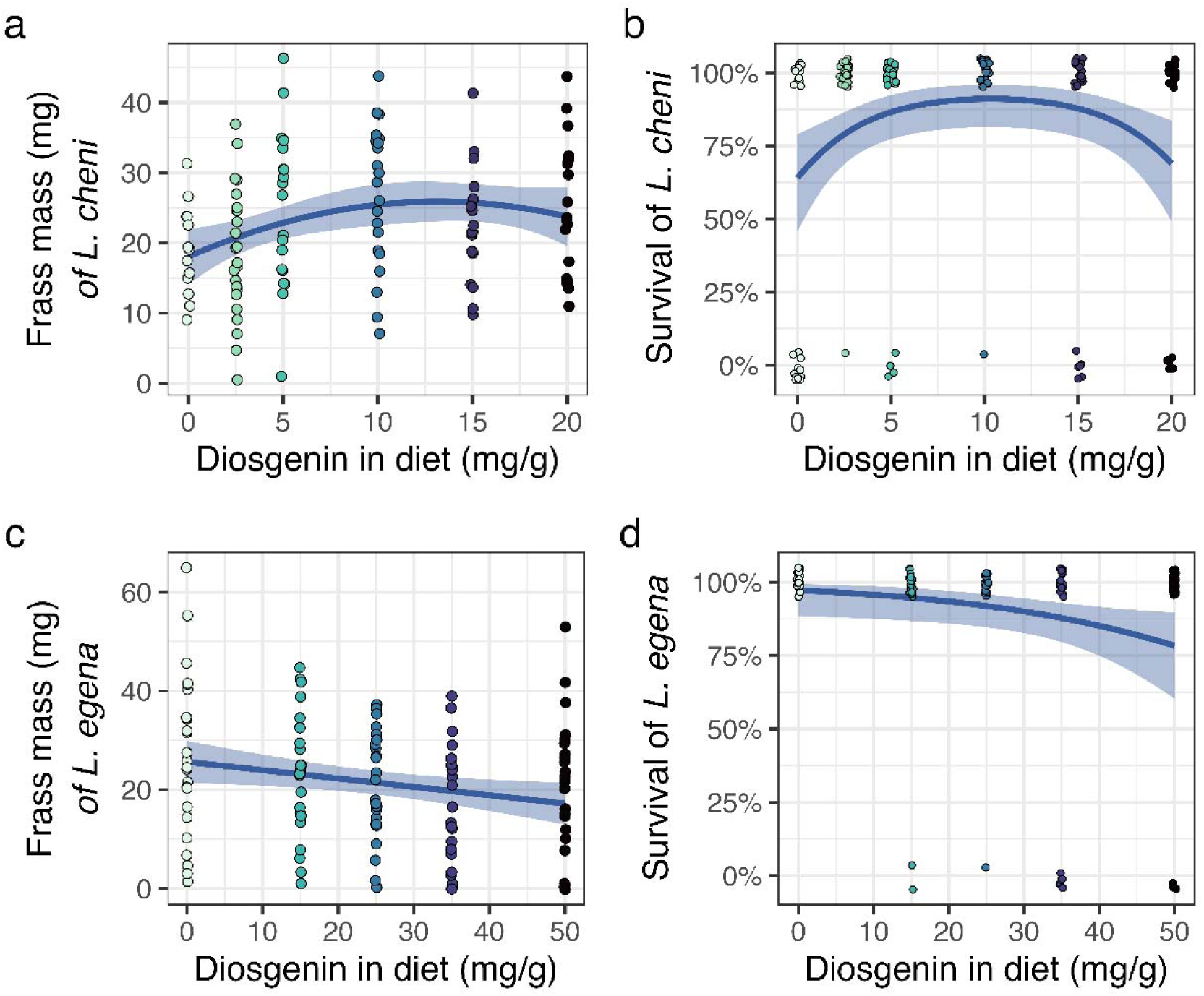
Effects of diosgenin concentration on feeding by *L. cheni* and *L. egena* beetles. Relationship between diosgenin concentration in the diet (mg/g) and (a) Frass weight (mg) of *L. cheni* beetles upon feeding (b) Survival rate (%) of *L. cheni* beetles (c) Frass weight (mg) of *L. egena* beetles upon feeding (d) Survival rate (%) of *L. egena* beetles. Each plot shows points for individual beetles and a fitted curve with shaded areas representing the 95% confidence intervals.

The quadratic model was also the best model to analyze the effect of diosgenin levels on the survival of *L. cheni* beetles, with the lowest AIC (141.48) compared to the linear model (AIC = 148.07) and the null model (AIC= 146.22). Like frass weight, survival of *L. cheni* increased initially at lower to intermediate concentrations of diosgenin (Slope = 0.339 ± 0.121 SE, z = 2.806, *p =* 0.005, Figure 3b), and declined thereafter at higher concentrations (Slope = −0.016 ± 0.006 SE, z = −2.802, *p =* 0.005, *R^2^* = 0.109, Figure 3b).

The evaluation of models to test the dose-dependent effects of saponins on the frass weight of *L. egena* as a response variable, against different doses of diosgenin as continuous predictor variable revealed that linear model has the lowest AIC (−694.06), although it was comparable to the quadratic model (AIC = −693.38), but both were lower than the null model (AIC= −690.39). The frass weight from *L. egena* decreased with increasing concentrations of the treatment (slope = −0.017 ± 0.007 SE, z = −2.408, *p* = 0.016, *R^2^* = 0.046, Figure 3c). To analyze the effect of diosgenin levels on the survival of *L. egena* beetles, the linear model had the lowest AIC (76.368), which was similar to the quadratic model (AIC = 76.815) but lower than and the null model (AIC= 80.020). Like frass weight, survival of *L. egena* declined with increasing concentrations of diosgenin (Slope = −4.564 ± 2.059 SE, z = −2.216*, p =* 0.023, *R^2^* = 0.052, Figure 3d).

## Discussion

Managing invasive plant species through classical biocontrol presents challenges due to unpredictable interactions among introduced natural enemies and their responses to host plant quality. This study specifically addressed the interactions between two specialist herbivores, *L. cheni* and *L. egena*, feeding on the invasive air potato vine. The results showed that *L. cheni* feeding on leaf tissue did not influence subsequent feeding by *L. egena* on bulbils as *L. egena* did not show significant differences in feeding on bulbils derived from JA-treated, *L. cheni* damaged, SA-treated, or control plants (Figure 1a-b). However, the performance of *L. egena*, as measured by pupal and eclosing adult weight, was significantly greater in the SA treatment compared to the control treatment (Figure 1c-d). However, there were no differences between any of the other comparisons. This finding was corroborated by the positive correlations between insect performance (pupal weight) and proxies of feeding (Frass weight (Figure 2a-d) and wax weight) and that varied slightly among treatment groups. This study highlights the nuanced interactions between specialist herbivores and host plant defenses, revealing that while *L. egena*’s feeding on air potato bulbils remained unaffected by prior damage, its performance was significantly enhanced by salicylic acid treatment, underscoring the complex dynamics in classical biocontrol systems. Collectively, these findings emphasize the importance of understanding whether previous feeding by *L. cheni* influences the food quality and drives the performance of the subsequent bulbil feeder, *L. egena*. Our results suggest that both insects can feed on the plant without negatively impacting the other.

In the dose-dependent feeding experiment, *L. cheni* exhibited non-linear frass production (proxy of feeding) and survival responses to diosgenin concentrations, with initial increases at lower to intermediate concentrations followed by negative effects at higher concentrations (Figure 3a-b). *Lilioceris egena* displayed a continuous decrease in frass production and survival with increasing diosgenin concentration (Figure 3c-d). Although, there is limited understanding of how these beetles interact under natural conditions in Florida, our findings contribute early and novel insights to the understanding of their interactions on the same host plant air potato and to the dose-dependent responses of these natural enemies to plant defense chemicals, specifically saponins. Overall, our findings best support the cumulative stress model, as both the beetles can inflict damage in an additive way, without negatively impacting each other’s feeding or performance. This is likely due to their niche differentiation on the plant, as they primarily feed on different plant structures, which also share different phytochemistry (Ghosh *et al*., 2013; Szakiel *et al*., 2017; Badole & Dighe, 2019; Kuncari, 2023; Sharma *et al*., 2024). These insights will improve understanding the possible effects of such interactions on biocontrol efficacy.

### Bulbil feeding experiment- *L. egena*: Plant-mediated indirect effects of *L. cheni* on L. egena

In the experiment where we tested the effect of different treatments on the feeding behavior and performance of *L. egena*, we expected that the larvae would exhibit increased feeding on bulbils originating from JA-treated plants and those derived from plants previously damaged by *L. cheni* herbivory, compared to control and SA-treated plants. This hypothesis is based on the previous findings that *L. cheni* herbivory on the foliage of air potato plants induced chemical defenses (i.e. Saponins) that acted as a feeding stimulant for *L. cheni* (Kaur *et al*., 2023). Thus, if induction of saponins is systemic and increases in the bulbils when leaves are damaged by *L. cheni*, subsequent feeding by *L. egena* may also increase. However, we found no difference in feeding damage by *L. egena* on the bulbils harvested from the four treatments (Figure 1a-b). Interestingly, we found that *L. egena* developed better on bulbils from SA-induced plants, as the pupae and subsequently emerging beetles gained more weight on bulbils from SA-induced plants compared to controls (Figure 1c-d). This contrasts with the performance of *L. cheni* on JA-induced or on prior herbivory feeding induced plants, where insects performed lowest on SA treated plants (Kaur *et al*., 2023). Additionally, we also found positive relationship between insect performance and feeding as a measure of food quality, which suggests that different treatments modulate this effect at different magnitudes (Figure 2a-d). This suggests that insect feeding on plants with varying defense qualities is closely linked to their performance. However, we found no effect of JA on the association between the performance and feeding, which diminishes the role of this phytohormone in this system.

One possible explanation for the difference observed in the previous results for *L. cheni* compared to the current findings for *L. egena* could be the nutritional richness of the bulbils, which are sources of sugars and other carbohydrates. Therefore, the defense chemistry of bulbils may differ significantly from that of the plant’s leaves, despite containing dioscin and diosgenin (Obidiegwu *et al*., 2020; Kuncari, 2023). For instance, studies have reported that *D. bulbifera* leaves are rich in phenols, tannins, alkaloids (Lin & Yang, 2008; Ghosh *et al*., 2013). Yams, closely related species in the genus *Dioscorea*, are known to be rich in carbohydrates, crude fat, crude fiber, and vitamin C (Ghosh *et al*., 2013; Ikiriza *et al*., 2022). Therefore, as the bulbils are more nutritious than leaves, the feeding preferences of *L. egena* likely respond more strongly to the nutritional aspects of the bulbils rather than their secondary chemistry. Additionally, we found no association between the performance and feeding in the JA treatment, which also does potentially hint that induced responses triggered by the JA pathway (e.g., saponins) could reduce *L. egena* feeding or performance (Figure 2c). Consistent with the findings from the dose-dependent experiment, it is likely that *L. egena* is negatively affected by saponins. This may be explained by its evolutionary adaptation to feed on bulbils, potentially increasing its susceptibility to saponins as an evolutionary trade-off. Also, since beetles performed significantly better in SA treatment (Figure 1c-d), it is likely that SA treatment in bulbils may not be linked to saponin production.

### Dose dependent bioassay - *L. egena* and *L. cheni –* feeding behavior and ecological specialization

The results from the dose-dependent assays also demonstrate distinct feeding strategies between *L. cheni* and *L. egena*, which have implications for their ecological roles and evolutionary trajectories*. Lilioceris cheni* followed the performance and feeding patterns of a sequestering specialist herbivore insect, while *L. egena* displayed the pattern of a generalist or non-sequestering specialist on air potato (Figure 3). Particularly, *L. cheni* fed (measured as frass mass) and performed (pupal weight and survival) better at low to intermediate saponin levels and declined at higher doses of diosgenin (Figure 3a-b). Therefore, the results for *L. cheni* are consistent with patterns predicted for a sequestering specialist (Karban & Agrawal, 2002; Ali & Agrawal, 2012). Previous literature suggests that sequestering specialists often show adaptations that allow them to handle specific phytochemicals efficiently up to a certain threshold beyond which their performance declines (Karban & Agrawal, 2002; Ali & Agrawal, 2012; Agrawal *et al*., 2021; Beran & Petschenka, 2022). Additionally, insects that sequester plant toxins are often better protected from predation and signal this protection with bright colors (Ernst 2005). Therefore, because *L. cheni* is highly specialized on *D. bulbifera* and is brightly colored aposematic beetle (Ernst, 2005), it is likely that it has evolved to sequester secondary metabolites, specifically dioscin and diosgenin that may also protect them from predators.

In contrast, the feeding behavior of *L. egena* aligns with predictions for a generalist or non-sequestering specialist, because its feeding and performance tended to decrease with increasing levels of saponins at high saponin levels (Figure 3c-d). This indicates a possibly different evolutionary path or adaptive strategy to plant defenses in *L. egena*, emphasizing less role of chemical sequestration strategies and more avoidance or tolerance as specialist behavior. A study also found similar patterns, reporting that non-sequestering specialists might employ strategies like rapid detoxification or avoidance of high-toxin areas of a plant (Hartmann, 2004; Erb & Robert, 2016). It is important to note that this study was conducted with freshly emerged adults, it would be interesting to test the dose dependent responses of larvae of both *L. cheni* and *L. egena* using artificial diet. However, a major limitation was that, in initial trial, the freshly emerged larvae did not feed well on the artificial diet, making answering that question challenging. Nevertheless, these results are broadly consistent with the results of the bulbil feeding assay, which used larval *L. egena*. Understanding feeding differences between *L. cheni* and *L. egena* is crucial for comprehending the niche differentiation and coexistence of these natural enemies on the same host plant.

### Ecological and evolutionary implications

The findings from this study underscore the complexity of natural enemy interactions with their host plants, highlighting how ecological pressures can drive divergent evolutionary paths even among closely related species. The differential responses to plant defense chemicals suggest that *L. cheni* and *L. egena* may have evolved under different selective pressures, leading to distinct adaptive strategies. This scenario is supported by broader ecological theories, such as those proposed by Ehrlich & Raven (1964), which posit that herbivore speciation and specialization are often closely tied to plant chemical defenses. In fact, a recent study suggested that repeated evolution of specific traits (like toxin resistance and sequestration) among herbivores across different taxonomic orders supports the predictability and repeatability of evolutionary outcomes under similar ecological pressures (Agrawal *et al*., 2024). In our system, the leaf feeder, *L. cheni*, appears to have evolved greater tolerance to saponins, whereas the bulbil feeder, *L. egena*, appears more susceptible to these saponins. The consistent differences in nutritional quality between leaves and bulbils likely is shaping how these species have adapted to the phytochemicals.

By integrating multiple natural enemies that target various aspects of the lifecycle of an invasive plant or its different plant structures, CBC can potentially leverage cumulative effects to enhance overall control. This strategy aligns with the cumulative stress hypothesis, which was supported in our system with two beetles that have evolved different feeding niches (leaves vs. bulbils). However, the dynamics among different natural enemies and their effects on host plants can vary, necessitating careful selection and management to optimize biocontrol efficacy. Understanding these complex interactions is crucial for developing robust biocontrol programs that can effectively mitigate the impacts of invasive species on biodiversity and ecosystem functions.

## Acknowledgements

We thank FDACS-DPI staff, particularly Rosemary Murray, for providing greenhouse space, bulbils, and the insects for carrying out the study.

## Statements & Declarations

### Author Contributions

JK, EK, and PGH conceived the research; JK and PGH designed the study; EK, OM, and PGH provided resources, JK and LN collected data; JK and PGH analyzed the data; JK wrote the first draft of the manuscript. All authors contributed to editing. All authors have read and agreed to the submitted version of the manuscript.

### Data availability statement

Data are available on Github Digital Repository (https://github.com/Plant-herbivory-interaction-lab/AirPotato_BulbilDoseDependent.git) and will be archived upon acceptance.□

