## Supplementary information for "Feeding niches drive specialist herbivore responses of conspecific insects to plant defenses in biocontrol"

Table S1.

| Ingredients | Diosgenin Concentrations (mg/g) | | | | | |
| --- | --- | --- | --- | --- | --- | --- |
| Diosgenin | 0 | 2.5 | 5 | 10 | 15 | 20 |
| Water (ml) | 60 | 60 | 60 | 60 | 60 | 60 |
| Colorado potato beetle mix (g) | 10 | 10 | 10 | 10 | 10 | 10 |
| Leaf material (g) | 0.4 | 0.4 | 0.4 | 0.4 | 0.4 | 0.4 |
| Agar (g) | 0.5 | 0.5 | 0.5 | 0.5 | 0.5 | 0.5 |
| KOH (g) | 0.003 | 0.003 | 0.003 | 0.003 | 0.003 | 0.003 |
| Neomycin (g) | 0.05 | 0.05 | 0.05 | 0.05 | 0.05 | 0.05 |

Table S2.

| Ingredients | Diosgenin Concentrations (g) | | | | |
| --- | --- | --- | --- | --- | --- |
| Diosgenin | 0 | 0.015 | 0.025 | 0.035 | 0.05 |
| Water (ml) | 60 | 60 | 60 | 60 | 60 |
| Colorado potato beetle mix (g) | 10 | 10 | 10 | 10 | 10 |
| Bulbil powder (g) | 0.4 | 0.4 | 0.4 | 0.4 | 0.4 |
| Agar (g) | 0.5 | 0.5 | 0.5 | 0.5 | 0.5 |
| KOH (g) | 0.003 | 0.003 | 0.003 | 0.003 | 0.003 |
| Neomycin (g) | 0.05 | 0.05 | 0.05 | 0.05 | 0.05 |
